# Siderophore synthetase-receptor gene coevolution reveals habitat- and pathogen-specific bacterial iron interaction networks

**DOI:** 10.1101/2023.11.05.565711

**Authors:** Shaohua Gu, Zhengying Shao, Zeyang Qu, Shenyue Zhu, Yuanzhe Shao, Di Zhang, Richard Allen, Ruolin He, Jiqi Shao, Guanyue Xiong, Alexandre Jousset, Ville-Petri Friman, Zhong Wei, Rolf Kümmerli, Zhiyuan Li

## Abstract

Predicting bacterial social interactions from genome sequences is notoriously difficult. Here, we developed bioinformatic tools to predict whether secreted iron-scavenging siderophores stimulate or inhibit the growth of community members. Siderophores are chemically diverse and can be stimulatory or inhibitory depending on whether bacteria possess or lack corresponding uptake receptors. We focused on 1928 representative *Pseudomonas* genomes and developed a co-evolution algorithm to match all encoded siderophore synthetases to corresponding receptor gene groups with >90% accuracy based on experimental validation. We derived community-level iron interaction networks to show that selection for siderophore-mediated interactions differs across habitats and lifestyles. Specifically, dense networks of siderophore sharing and competition were observed among environmental (soil/water/plant) strains and non-pathogenic species, while only fragmented networks occurred among human-derived strains and pathogenic species. Altogether, our sequence-to-ecology approach empowers the analyses of social interactions among thousands of bacterial strains and uncovers ways for targeted intervention to microbial communities.

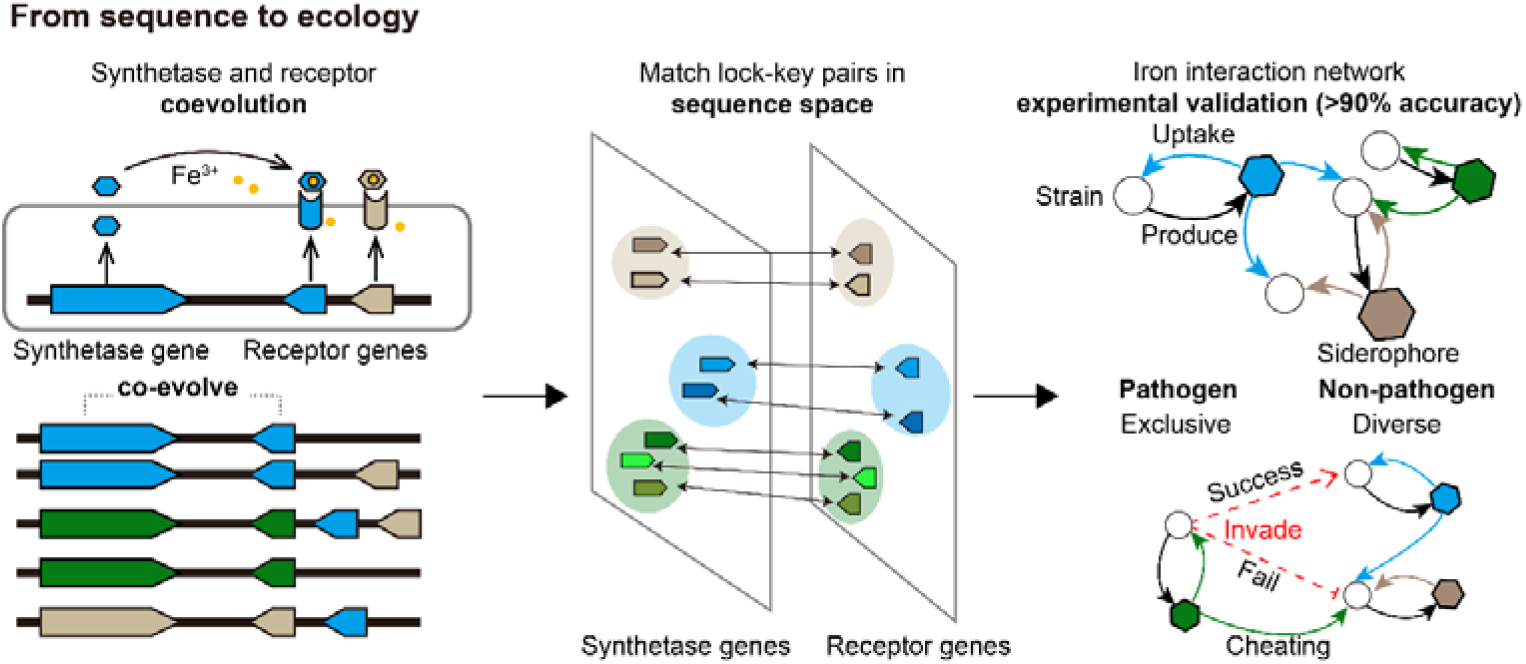

## Introduction

Microbial communities populate all ecosystems on earth from terrestrial to aquatic environments, impacting human health, agriculture, and industry ^1-3^. The dynamics and functioning of these communities are shaped by complex and unexplored interactions between microorganisms ^4,5^. As the number of sequenced microbial genomes continues to grow ^6,7^, there is enormous interest in developing approaches to predict microbial interaction networks based on genomic data. Such efforts are essential to obtain complete insights into community functioning as many microorganisms cannot be cultured in the laboratory ^8^, while their ecological roles could still be inferred through sequence-to-interaction mapping. Currently, sequence-to-interaction mapping approaches primarily focus on metabolic interactions, with Genome-scale Metabolic Models (GEMs) serving as the primary tool for establishing the pan-reactome of microbial communities ^9,10^. These methods infer metabolic reactions from the genome annotation of enzymes, and then reconstruct a flux model to understand how microorganisms take up essential nutrients and release metabolic byproducts into the environment ^11-13^.

Despite the significance of primary metabolism, there is increasing evidence that also other secreted compounds synthesized through secondary metabolism ^14^, play a major role in shaping microbial interactions ^15,16^. Nearly all microbes actively synthesize and secrete compounds to fulfill a diverse set of functions, including communication, resource scavenging, motility, and attack of and defense against competitors ^17^. Many of these secreted compounds were previously considered non-essential for microbial growth in laboratory settings, but have since been shown to be critical for competitiveness in natural environments ^15,16^. However, sequence-to-interaction mapping has rarely been applied to secreted compounds, particularly because the synthesis and mode of action of secondary metabolites is challenging to predict.

Here, we developed a secondary metabolite sequence-to-interaction approach focusing on iron-scavenging siderophores, one of the most prevalent and diverse classes of microbial secondary metabolites ^18^. Iron is critical for microbial growth and survival, because of its importance as catalytic group in enzymes guiding key biological processes such as respiration and DNA replication ^19^. However, the concentration of bioavailable iron is typically below the required level in most habitats ^19-21^, and upon iron limitation, nearly all bacteria produce siderophores that efficiently chelate iron from insoluble environmental stocks ^22,23^. Siderophores are typically diffusible and able to chelate iron over a broad physical range ^24^. Once iron is bound, the complex is recognized and taken up by specific receptors embedded into the bacterial cell membrane ^23^. Diffusible siderophores mediate several types of social interactions. They can cooperatively be shared among contributing bacterial strains with matching receptors for the uptake of the iron-siderophore complex ^23,25^. Siderophore can also be deployed as competitive agents to limit access to iron if co-occurring strains lack matching receptors ^23,26^. Finally, siderophores can be exploited by cheaters bacteria that have receptors for siderophore uptake but do not pay the cost of producing siderophores themselves ^22,23^. Consequently, while siderophore-mediated interactions have important impacts on microbial community dynamics and functions ^27-30^, we still poorly understand how these interactions scale up at the network level in environmental and in host-associated bacterial populations.

The aim of our study was to infer how receptors and siderophores have co-evolved and use this information to develop algorithms that identify matching siderophore-receptor pairs that predict interaction networks in bacterial communities based on sequence data. We previously used the genome sequences of 1928 *Pseudomonas* strains to develop bioinformatic pipelines that allowed us to predict the chemical structure of 188 pyoverdines (the main siderophores of this genus) and to identify 4547 FpvA-receptor genes (segregating into 94 groups) involved in pyoverdine uptake ^31^. Here, we capitalize on this work to develop a Co-evolution Pairing Algorithm to match the pyoverdines (key) and receptors (lock) into 47 unique lock-key groups and over 90% of these predicted interactions could be validated experimentally *in vivo*. Using the predicted lock-key pairs, we then reconstructed siderophore-mediated iron interaction networks among all *Pseudomonas* strains. We found that network complexity was high among strains isolated from soil-, water-, and plant-derived habitats, whereas complexity was lower among strains isolated from human-associated habitats. We further noticed that interaction networks among pathogenic species were small and loose with few pyoverdine interactions existing between strains. The opposite was the case for strains from environmental habitats. Taken together, the developed sequence-to-interaction mapping tool can accurately predict social interaction networks mediated by siderophores in complex bacterial communities. Our findings suggest that selection for social interactions varies across habitats and lifestyles, thus providing novel insights into community functions and connectivity.

## Results

### Three classes of pyoverdine strategies in *Pseudomonas* strains and the lock-key (receptor-synthetase) principle of co-evolution

Our data set consists of 1928 nonredundant *Pseudomonas* strains, producing a total of 188 chemically different pyoverdine types and featuring 94 different receptor groups according to our recently developed bioinformatic prediction tools^31^. We first explored the diversity of strains in terms of phylogeny, ecological habitat, and pyoverdine strategies. At the phylogenetic level, our data set included a diverse set of *Pseudomonas* species, where *P. aeruginosa* (28.7%), *P. fluorescens* (7.0%), *P. syringae* (6.0%), and *P. putida* (2.2%) were the most abundant ones (Figure 1a). The strains originated from diverse habitats, including human-derived habitats (21.2%), soils (13.6%), plants (12.1%), and water (6.4%), although the origin of many strains (39.5%) is unknown (Figure 1a).

**Figure 1.**
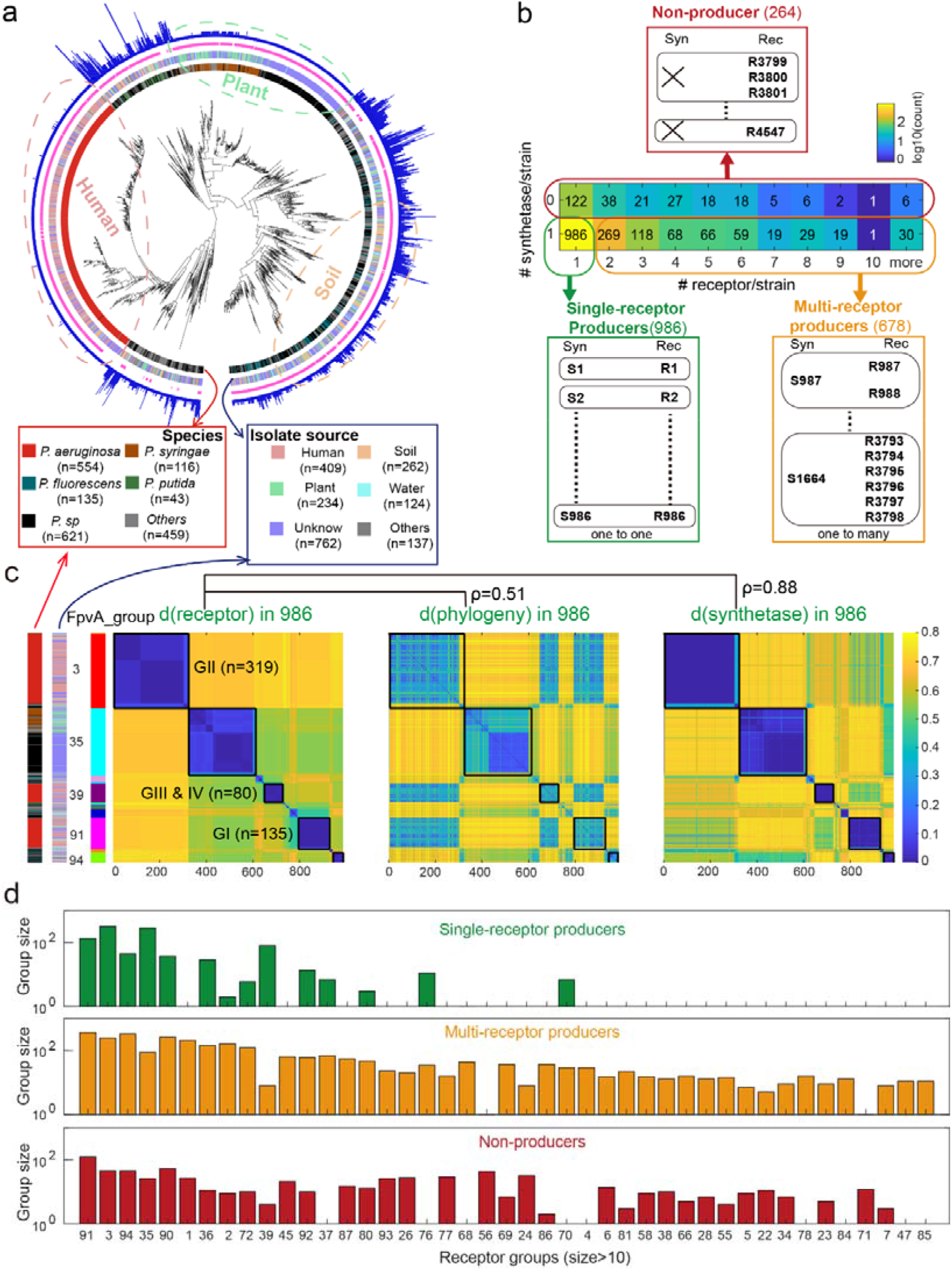
Classification of *Pseudomonas* strains and elucidation of the co-evolution between pyoverdine synthetases and receptors. **a.** Phylogenetic relationship among the 1928 *Pseudomonas* strain based on the concatenated alignment of 400 single-copy conserved genes. Starting from inside, colors in the first ring distinguish the five most prevalent species, with “Others” representing the remaining less abundant species. Colors in the second ring distinguish the four most prevalent sources of isolation. In the third ring, claret and blank regions cover strains with complete pyoverdine synthetase clusters and strains without synthetase gene clusters, respectively. In the fourth blue ring, the bar height indicates the number of FpvA receptors present in each strain. **b.** Strains can be classified into three types by scoring the presence/absence of a synthetase cluster and counting the number receptors in each genome: (i) single-receptor producers containing one pyoverdine synthetase cluster and one FpvA receptor gene; (ii) multi-receptor producers containing one pyoverdine synthetase cluster and several FpvA receptor genes; and (iii) Non-producers lacking synthetase gene but containing at least one receptor gene. **c.** Heatmap visualizing distances between feature sequences of the FpvA receptors and the pyoverdine synthetase clusters and between FpvA features sequences and phylogenetic genes among the 986 single-receptor producers. In all three heatmaps, the hierarchical clustering of the strains follows the one used for the FpvA feature sequences (left panel). The black squares on the heatmaps denote the five major FpvA groups. Three of these groups correspond to the receptors found among *P. aeruginosa* strains and are labelled with black text. **d.** The 43 largest FpvA receptor groups with more than 10 members (sorted by group size) and their frequency among single-receptor producers, multi-receptor producers, and non-producers.

To assess the diversity of pyoverdine production and uptake strategies, we analyzed the absence or presence of pyoverdine synthesis clusters and counted the number of FpvA pyoverdine receptors per strain. We found three types of pyoverdine-utilization strategies (Figure 1b). “Single-receptor producers” were the most common type (985 strains, 51.1%) and refer to strains with one pyoverdine synthesis locus and one FpvA receptor gene. “Multi-receptor producers” were the second most common type (679 strains, 35.2%) and includes strains with one pyoverdine synthesis cluster but multiple FpvA receptor genes. “Non-producers” were the least common type (264 strains, 13.7%) and refer to strains that lack the pyoverdine synthesis cluster but contain at least one receptor gene. While strains can possess multiple FpvA receptor genes, no strain carried more than one pyoverdine synthesis cluster. This observation is in line with the expected high costs of pyoverdine synthesis, which is based on a series of gigantic modular enzymes known as NPRS (non-ribosomal peptide synthetases)^32^.

Based on these findings, we hypothesized that in each single-receptor producer, the sole receptor present should recognize the self-produced pyoverdine to ensure fitness benefits when faced with iron limitations. Consequently, synthetase and receptor pairs should reflect molecular co-evolution, where mutational alterations in the synthetase structure should select for corresponding changes in the receptor sequences to preserve the lock-key relationship and efficient iron uptake. To test this hypothesis, we focused on the 986 single-receptor producers and calculated the degree of covariation between sequence distances matrices of the receptor, the synthesis cluster, and 400 conserved genes. For the receptors (FpvA) and the synthesis cluster, we used the feature sequences that are most predictive of receptor specificity and pyoverdine molecular structure, as identified in our previous work^31^. In support of the co-evolution hypothesis, we found a strong correlation between the distance matrixes of the receptors and the synthesis clusters (Pearson’s r=0.88), a correlation that is much stronger than between the receptor and the phylogeny matrix (Pearson’s r=0.51) (Figure 1c). Notably, we observed strong clustering patterns in the sequence space of the receptors, forming distinctive blocks that closely match with the clustering patterns of their corresponding pyoverdine synthesis clusters. Using our receptor clustering pipeline^31^, we identified 17 receptor groups among the 986 single-receptor producers. Importantly, three out of the 17 receptor groups represent the FpvA receptors found in the human pathogen *P. aeruginosa* (labelled as GI, GII and GIII+IV in Figure 1c left panel) for which the selective uptake of the corresponding pyoverdines has been demonstrated^22,33^. These analyses strongly indicate that cognate receptors and synthesis genes have co-evolved in single-receptor producers, resulting in one-to-one “lock-key” relationships.

Co-evolutionary lock-key groups cannot directly be inferred for multi-receptor producers because there is currently no method to distinguish the “self-receptor” responsible for absorbing the self-produced pyoverdine from the other FpvA receptors responsible for the uptake of heterologous pyoverdines produced by other strains. Moreover, receptor diversity seems to be much larger among multi-receptor than among single-receptor producers. Indeed, when focusing on the 43 (out of the 94) receptor groups with more than 10 members, we found that single-receptor producers covered only 14 of these groups (32.6%), while multi-receptor producers had a much more diverse receptor coverage (41 groups, 95.3%) (Figure 1d). Non-producers had a similarly broad receptor coverage (34 groups, 79.1%). We also found that single-receptor producers tend to connect more compactly (mean silhouette index = 0.96±0.16) than multi-receptor producers (mean silhouette index = 0.78±0.19) and non-producers (mean silhouette index = 0.79±0.20) in the sequence space (Figure S1 and S2), suggesting that receptors from single-receptor producers might evolutionarily be more conserved than receptors from non-producers and multi-receptor producers. Altogether, these observations imply that single- and multi-receptor producers take on different roles in iron interaction networks.

### Development of the Co-evolution Pairing Algorithm and its experimental validation to predict iron interaction networks

The aim of this section is to establish a lock-key receptor-pyoverdine interaction map across all three strain types. A first task in this process is to identify receptors in multi-receptor producers that are used to take up the self-produced pyoverdine. A first intuitive approach was to check for receptors proximate to the pyoverdine synthetase genes (Solution 1), while an alternative approach was to use the lock-key pairs identified for single-receptor producers and analysis whether similar pairs occur in multi-receptor strains (Solution 2). Even after completing these two solutions, more than half of the receptor groups could not be paired with any pyoverdine synthetases. Specifically, solution 1 identifies putative self-receptors in 87.1% (591 out of 678) of the multi-receptor producers, while the solution 2 could be applied to only 68.7% (466 of 678) of multi-receptor strains.

We thus developed an unsupervised learning algorithm, termed “Co-evolution Pairing Algorithm” (Solution 3), which searches for the set of synthetase-receptor combinations that maximizes co-evolutionary association based on feature sequence distance between synthetase and receptor (Figure 2a-b, see Method 4 for details). We then checked for consistency in self-receptor identification across the three solutions (Figure 2c). Solution 1 and Solution 2 yield high levels of consistency (99.5% across 433 strains). The unsupervised Solution 3 shares high consistency with Solution 1 (93.7%, across 591 strains) and Solution 2 (94.4%, across 466 strains), indicating that all three solutions are legitimate, with Solution 3 having the advantage of being applicable to all strains.

**Figure 2.**
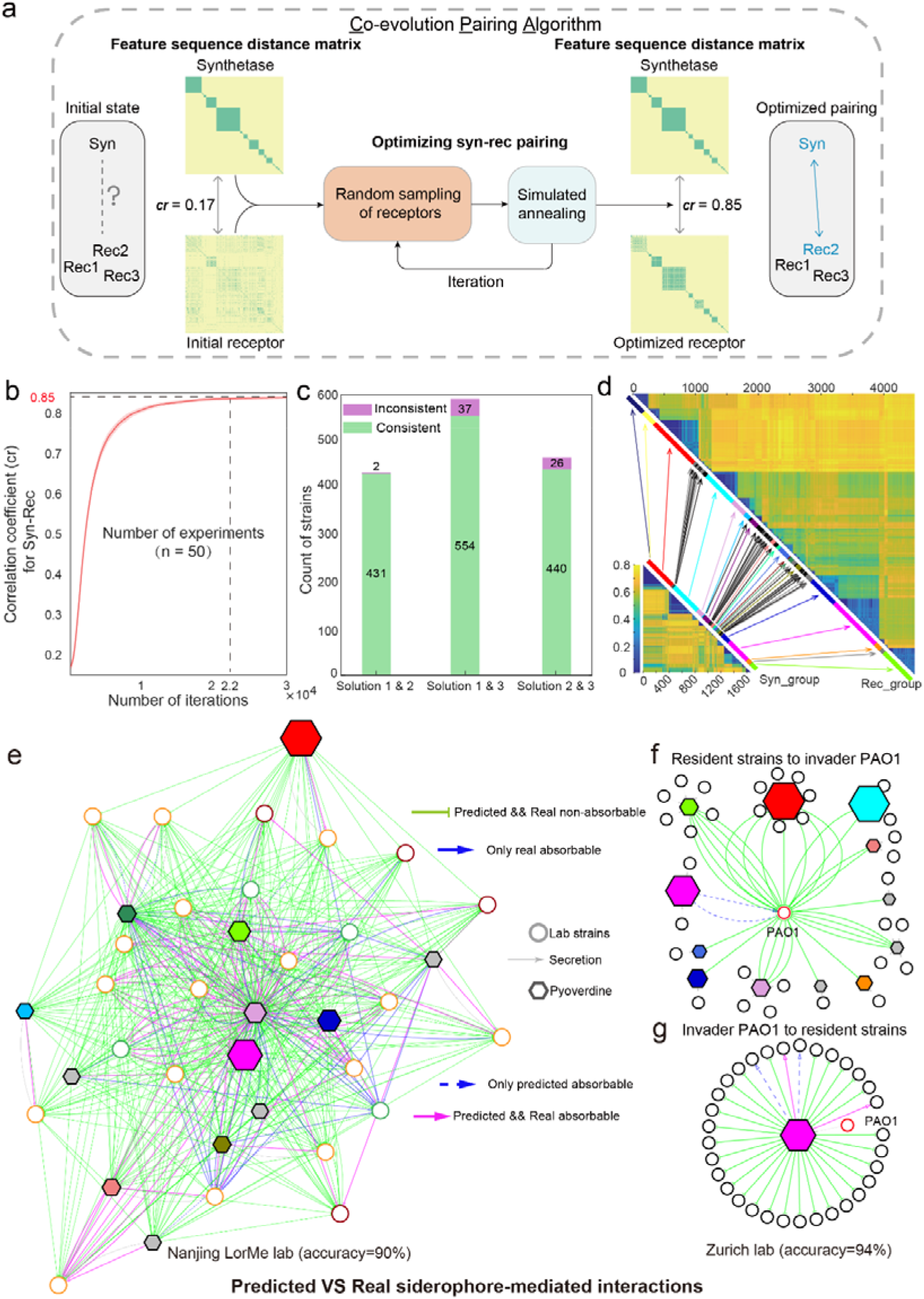
Unsupervised co-evolutionary algorithm to establish a lock-key pair map for pyoverdine synthetase and receptor groups and its experimental validation. **a.** Flowchart depicting the Co-evolution Pairing Algorithm (Solution 3) to match the synthetase in each strain to its “self-receptor”, based on an unsupervised learning scheme that optimizes co-evolutionary strength (correlation coefficients) between the feature sequence distance matrices of synthetases and matched receptors. The mean correlation coefficients (cr) between the two matrices before and after the optimization process are shown. **b.** The correlation coefficient (cr) and stability of the algorithm were examined by multiple rounds of learning (exp 1 to 50). **c.** Consistency of identifying the same receptor as self-receptor across three different solutions. Solution 1: receptor is within 20,000 bp proximity to the synthetase. Solution 2: the lock-key pairs of single-receptor producers are applied to multi-receptor producers. Solution 3: unsupervised co-evolution pairing algorithm. **d.** Predicted lock-key pairs connected in sequence space: 1664 synthases (bottom left) linked to the 4547 receptors (top right). Arrows depict the 47 lock-key links between synthetase and receptor groups. The colored (use the same color code as Figure 1c) and black/ grey shaded lines represent groups with single-receptor producers and without single-receptor producers, respectively. **e.** Predicted vs. observed iron interaction network among the 24 experimental strains. Each circular node represents an experimental strain and line colors stand for single-receptor producers (green), multi-receptor producers (yellow) and non-producers (red). Hexagons represent the predicted 13 lock-key receptor-pyoverdine groups. Edges from strain nodes to lock-key nodes represent pyoverdine production, while edges from lock-key nodes to strain nodes represent utilization. Green (non-usable pyoverdine) and pink (usable pyoverdine) edges depict cases in which experimental observations match bioinformatically predicted interactions. Blue edges depict incorrectly predicted pyoverdine interactions. The pyoverdine groups that are produced by at least one single-receptor producer are shown as colored hexagones with the color of the respective receptor group (Fig. 1c), whereas grey hexagons depict the pyoverdine groups that are exclusively produced by multi-receptors. **f+g.** Predicted vs. observed iron interaction networks based on data from a previous study carried out in the Zurich lab. The predicted interactions were inferred by the algorithms presented in this study, while the experimental data is taken from Table S2 of Figueiredo et al. ^34^.

Our Co-evolution Pairing Algorithm, allocated the self-receptors of the 1664 pyoverdine-producing strain into 47 distinct lock-key groups based on the receptor feature distance (Figure 2d and Figure S3). Most self-receptors belonged to 17 lock-key groups (single-receptor producers: 986 = 100%, multi-receptor producers: 572 = 84.4%), while the remaining self-receptors of multi-receptor producers (106 = 15.6%) segregated into 30 additional receptor groups (Figure S3). Out of the total 4547 FpvA genes detected, we identified 2883 receptors that are not self-receptors and thus possibly serve as “cheating receptors” to take up heterologous pyoverdines produced by other strains. Most of these cheating receptors (2703 = 93.8%) also segregated into the 47 lock-key groups, confirming that they could be used to exploit at least one of the 188 produced pyoverdines. The remaining cheating receptors (180 = 6.2%) could not be linked to any of the 47 lock-key groups, suggesting the existence of rare receptor groups that presumably match rare pyoverdine structures not covered by our dataset (Figure S3).

By combining co-evolution pairing algorithm and the lock-key pairing, we predicted siderophore-mediated iron interaction networks, based on pyoverdines that were produced and could be taken up by corresponding receptors by community members. We conducted two experiments to validate predicted interactions. For the first validation, we used a *Pseudomonas* community from the Laboratory of rhizosphere microbial ecology (LorMe) in Nanjing (China), which was originally isolated from the tomato rhizosphere ^29^. We included 24 independent strains and subjected their genomes to our bioinformatic pipelines to predict pyoverdine structures, to find all FpvA receptors^31^, to identify self-receptors, and to allocate pyoverdines and receptors into lock-key groups. We found that these 24 strains included 4 single-receptor producers, 16 multi-receptor producers and 4 non-producers (Figure S4) and that their self-receptors could be allocated to 13 lock-key groups (Figure S5). With this information, we predicted the pyoverdine-mediated iron interaction network between the 24 strains (Figure 2e). For the experimentally validation, we confirmed the pyoverdine production status of the 20 putative producers and the 4 non-producers (Figure S4). We then followed a modified version of our previously established protocols to calculate the net effect pyoverdine has on the growth of other strains (GE_Pyo_), while controlling for the effects of other metabolites in the supernatant ^29^. This approach allowed us to obtain an experimentally derived pyoverdine-mediated iron interaction network (Figure 2e and Figure S6). We found that 90% of the observed interactions (whether strains are stimulated or inhibited by the pyoverdines of others) matched the computationally predicted interactions from sequence data.

The second experimental validation involved strains from the Zurich (Switzerland) collection, isolated from soil and freshwater habitats ^23^. In this case, we used published experimental data from the literature ^34^. The focus of this earlier study was to test whether the opportunistic human pathogen *P. aeruginosa* PAO1 can invade natural soil and pond communities based on its ability to use pyoverdine produced by the natural isolates. We used data from all the strains for which genome sequences were available (PAO1 and 33 natural isolates), to establish pyoverdine-mediated iron interaction networks (Figure 2f-g). We then applied our bioinformatic pipelines as explained for the Nanjing collection and found a high level of consistency (94%) between the predicted and observed pyoverdine-mediated iron interactions (Figure 2f-g). Altogether, the two validation experiments demonstrate that siderophore-mediated microbial interactions can accurately be predicted based on genome-sequence analysis using the lock-key relationship between receptor and synthetase genes.

### Pyoverdine-mediated iron interaction networks vary across habitats

Following the successful validation, we applied the lock-key pairing methodology to our full data set to construct the pyoverdine-mediated iron interaction network among all 1928 *Pseudomonas* strains (Figure 3a). To keep traceability in such an enormous network, we allocated strains into microbial siderophore functional groups, defined as strains that produce the same pyoverdine types and can utilize the same repertoire of pyoverdines. Overall, the network featured 407 microbial siderophore functional groups, 47 different lock-key receptor-pyoverdine groups, 307 production edges and 1788 utilization edges.

**Figure 3.**
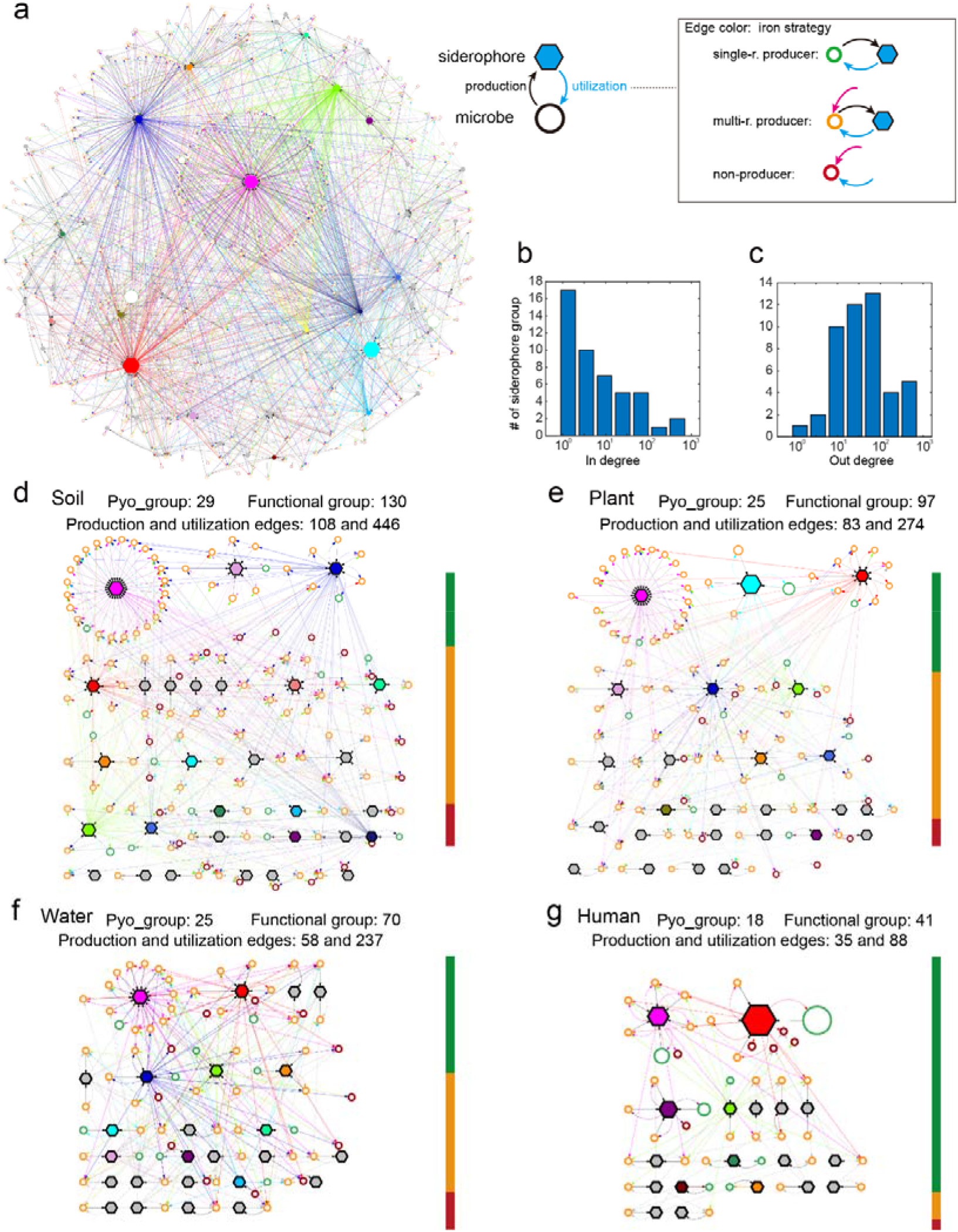
*Pseudomonas* iron interaction networks vary across habitats. **a.** The predicted iron interaction networks mediated by pyoverdines among 1928 *Pseudomonas* strains. Circular nodes represent functional groups (i.e., strains that produce the same pyoverdine type and utilize the same repertoire of pyoverdines) with node size being proportional to the number of strains within this functional group. Line colors of circular nodes represent single-receptor producers (green), multi-receptor producers (yellow) and non-producers (red). Hexagonal nodes represent lock-key pyoverdine groups with node size being proportional to the number of strains utilizing the corresponding pyoverdine. The hexagons of the 17 pyoverdine groups found among single-receptor producers are highlighted with the corresponding receptor group colors (as shown in Fig. 1c), while the pyoverdine groups that are exclusively found among multi-receptor producers are depicted by grey hexagons. Edges from circular to hexagonal nodes represent pyoverdine production, while edges from hexagonal to circular nodes represent pyoverdine utilization (with edge color matching the color of the functional group). **b.** The in- and **c.** the out-degree distribution of the pyoverdine nodes. The in- and out- degrees are defined by the number of edges pointing towards (representing production) or originating from (representing utilization) a pyoverdine node. **d-g.** Iron interaction networks of strains isolated from soil, plant, water and human habitats. The color bars on the right of each panel show the proportion of single-receptor producers (green), multi-receptor producers (yellow) and non-producers (red). Node symbols and colors are the same as in panel **a.**

Our iron interaction network can be considered as a special version of a bipartite network, characterized by two types of nodes (microbial functional groups and pyoverdine groups) and two types of directional edges (utilization and production). In ecological bipartite networks such as those associated with pollination and food webs, topology plays a crucial role for ecological functions and community assemblies ^35^. The topological metrics of our network reveals significant heterogeneity. Specifically, the in- and out-degree distribution of pyoverdine nodes is heavy-tailed, indicating “hub” siderophores that are either produced or utilized more extensively than others (Figure 3b-c). This high degree of heterogeneity suggests non-random network structure, likely influenced by specific eco-evolutionary forces (*e.g.,* fierce competition for iron). Indeed, we observed higher than expected yet moderate modularity (Qb=0.51, compared to Qb=0.41 in a randomized network) and nestedness (NODF=0.15, compared to NODF=0.04 in a randomized network). These values differ from mutualistic networks that typically feature high nestedness (*e.g.,* pollination networks), while antagonistic networks are typically characterized by high modularity (*e.g.,* herbivory and host-parasite networks) ^35,36^. Taken together, the iron interaction network may represent a novel type of ecological bipartite network and stands out as one of the largest reported to date.

Next, we created separate networks for strains isolated from soil (262 strains), plant (234), water (124), and human-derived (409) habitats. We found that frequencies of the three pyoverdine strategies and network topologies varied fundamentally between the four habitats (Figure 3d-g). For example, among the soil-derived strains, there were 56.9% multi-receptor producers, 27.5% single-receptor producers and 15.7% non-producers (Table S1). In contrast, there were only 10.0% multi-receptor producers and 4.0% non-producers, but 86.1% single-receptor producers among human-derived strains. Regarding network topologies, we observed that the number of microbial functional groups was higher for soil (130, value scaled to number of strains = 0.50), plant (97, 0.41), and water (70, 0.56) habitats than for human-related habitats (41, 0.10). Moreover, many functional groups (60.7%) exclusively occurred in a single habitat: soil (80, 23.7%), plant (56, 16.6%), water (43, 12.7%), and human (26, 7.7%), whereas only 8 functional groups (Figure S7a) and 11 pyoverdine groups (Figure S7b) were present in all four habitats. Altogether, these results indicate that the eco-evolutionary forces shaping the iron interaction networks differ substantially across habitats.

### Pyoverdine-mediated iron interaction networks vary between pathogenic and non-pathogenic species

*Pseudomonas* spp. do not only populate a variety of habitats but can also display diverse lifestyles. The most prominent division in lifestyles occur between pathogenic and non-pathogenic species. Here, we explore whether iron interaction networks differ between these two lifestyles. We allocated strains to pathogenic and non-pathogenic species groups and predicted the iron interaction networks for all species with more than five strains (Fig. 4a and Table S2). The most abundant pathogenic species were the human pathogen *P. aeruginosa* and the plant pathogen *P. syringae*, while the most abundant non-pathogenic environmental species were *P. fluorescens* and *P. putida*, of which many are neutral or even beneficial for hosts.

**Figure 4.**
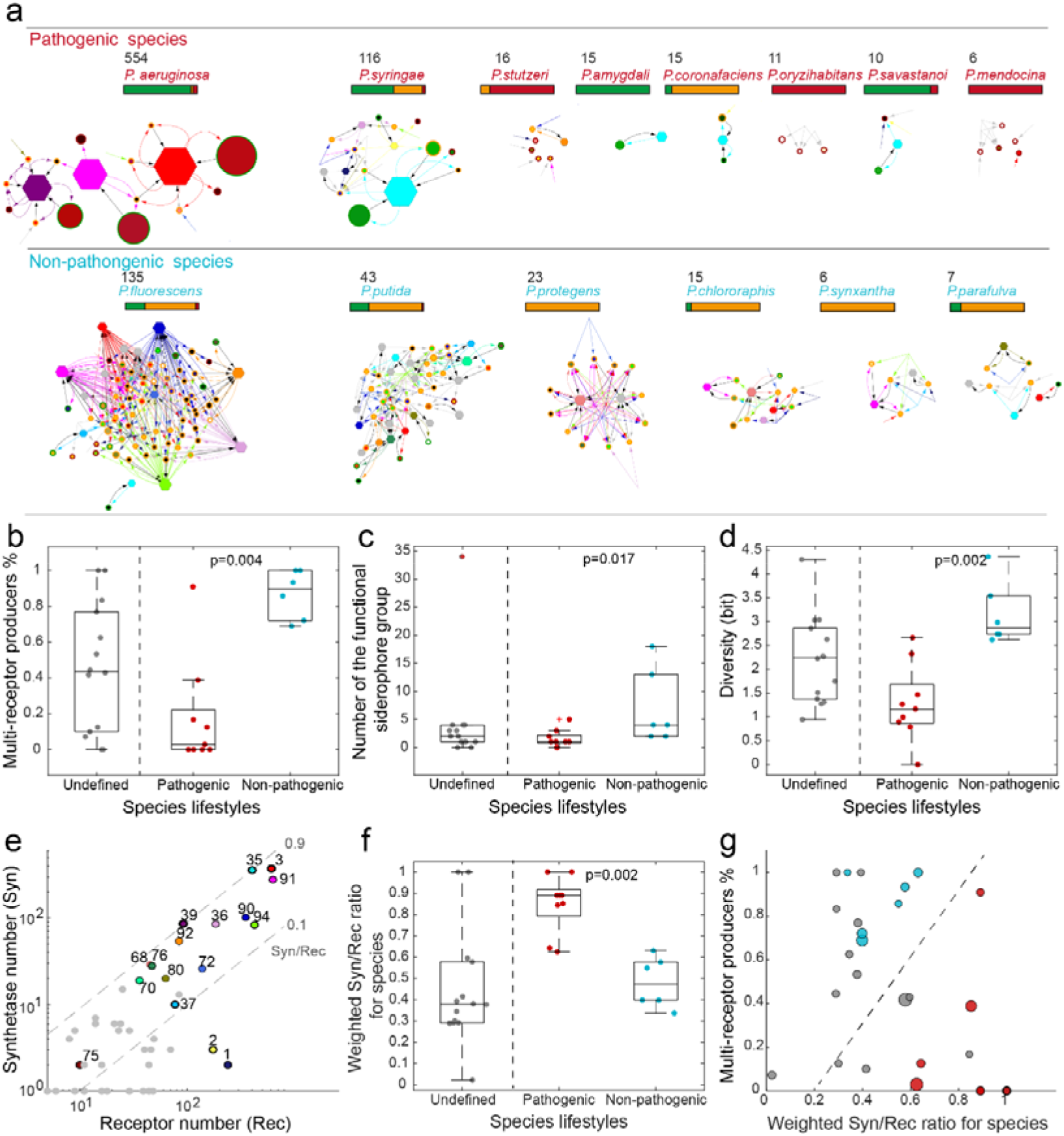
Network properties differ between pathogenic and non-pathogenic *Pseudomonas* spp. **a.** The pyoverdine-mediated iron interaction network for pathogenic (top row) and non-pathogenic (bottom row) species with more than five strains in our database. Symbols and color codes match the ones in Figure 3a. **b-c.** The difference of the percentage of multi-receptor producers and the number of the microbial functional group between different species lifestyles. Pathogenic species are colored red, non-pathogenic species are colored blue, and species that do not have clear role in pathogenicity are colored gray. **d.** The difference of diversity quantified by the entropy of their functional group frequencies between different species lifestyles. **e.** Counts of annotated receptors (x-axis) and annotated synthetase (y-axis) for each lock-key group. Lock-key groups are colored by as these in Figure 3a. The grey dotted line represents 0.1 and 0.9 ratios of the synthetase count verse receptor count (termed as “Syn/Rec”). **f.** The difference of the weighted Syn/Rec ratio for each species between different species lifestyles. **g.** The percentage of multi-receptor producers (y-axis) and the weighted Syn/Rec ratio (x-axis) for each species clearly separates pathogenic and non-pathogenic strains (segregation illustrated by the dashed line). The size of the dot represents its diversity quantified by the entropy of their functional group frequencies. Same color coding as that in **b**.

We observed multiple differences in network topologies between the two lifestyles (Figure 4a). First, strains of pathogenic species were mostly single-receptor producers or non-producers (Figure 4a-b), while strains of non-pathogenic species were primarily multi-receptor producers (Figure 4a-b). Second, the diversity of siderophore functional groups was much lower in pathogenic compared to non-pathogenic species (Figure 4c and Table S2). Consequently, the complexity of iron interaction networks (quantified by the entropy of their functional groups) was lower in pathogenic than in non-pathogenic species (Figure 4c). For example, *P. aeruginosa* (the most abundant species in our dataset, 554 strains), has a simple interaction network with only three functional groups, while *P. fluorescens* (135 strains) has a complex interaction network with 13 functional groups (Table S2). Finally, we calculate the Syn/Rec value defined as the ratio of synthetase and receptor groups present in a lock-key group (Figure 4e). In each lock-key group, we calculated the Syn/Rec ratio as the count of synthetase divided by the count of receptors. A Syn/Rec ratio near one indicates that there are no cheating receptors in this lock-key group, *i.e.*, this pyoverdine is exclusive to its producers and cannot be utilized by strains not producing it. Conversely, a Syn/Rec ratio near zero means that most receptors in this lock-key group are cheating receptors, and the corresponding pyoverdine is more sharable and exploitable.

We found that the Syn/Rec ratio was much lower in non-pathogenic than in pathogenic species (Figure 4f). Furthermore, Syn/Rec ratio differed substantially across lock-key groups (Figure 4e). For example, the lock-key groups 39 (purple hexagon, presents in *P. aeruginosa*) and 35 (cyan, presents in several plant pathogens, Fig. 4a) had high Syn/Rec ratios of 0.93 and 0.89, respectively. In contrast, the lock-key group 94 (light green, presents in *P. fluorescens*) had a much lower Syn/Rec ratio of 0.20. Collectively, the proportion of multi-receptor producers and the preference for producing shareable or exclusive siderophores clearly differentiated between pathogenic and non-pathogenic species on the two-dimensional plane (Figure 4g). Altogether, these results show that non-pathogenic species form open networks dominated by shareable pyoverdine lock-key groups, whereas pathogenic species form more closed networks dominated by exclusive pyoverdine lock-key groups.

### Modeling the relationship between pyoverdine utilization strategies and community dynamics

In order to explore the connection between pyoverdine utilization strategies of individual strains and the resulting community dynamics, we build simple ecological models of siderophore-mediated ecological competition (Figure 5a) ^37^. Our model revealed that different single-receptor producers cannot co-exist, illustrating the competitive exclusion principle ^37^ and matching our observation that a high proportion of single-receptor producers were associated with simple network structures (Figure 4a). In contrast, multi-receptor producers were more likely to coexist based on our model (Figure 5b), mirroring our observation that a high proportion of multi-receptor producers correlated with increased network diversity and complexity (Figure 3 and Figure 4a). Finally, non-producers could not exist in the absence of producers in our models.

**Figure 5.**
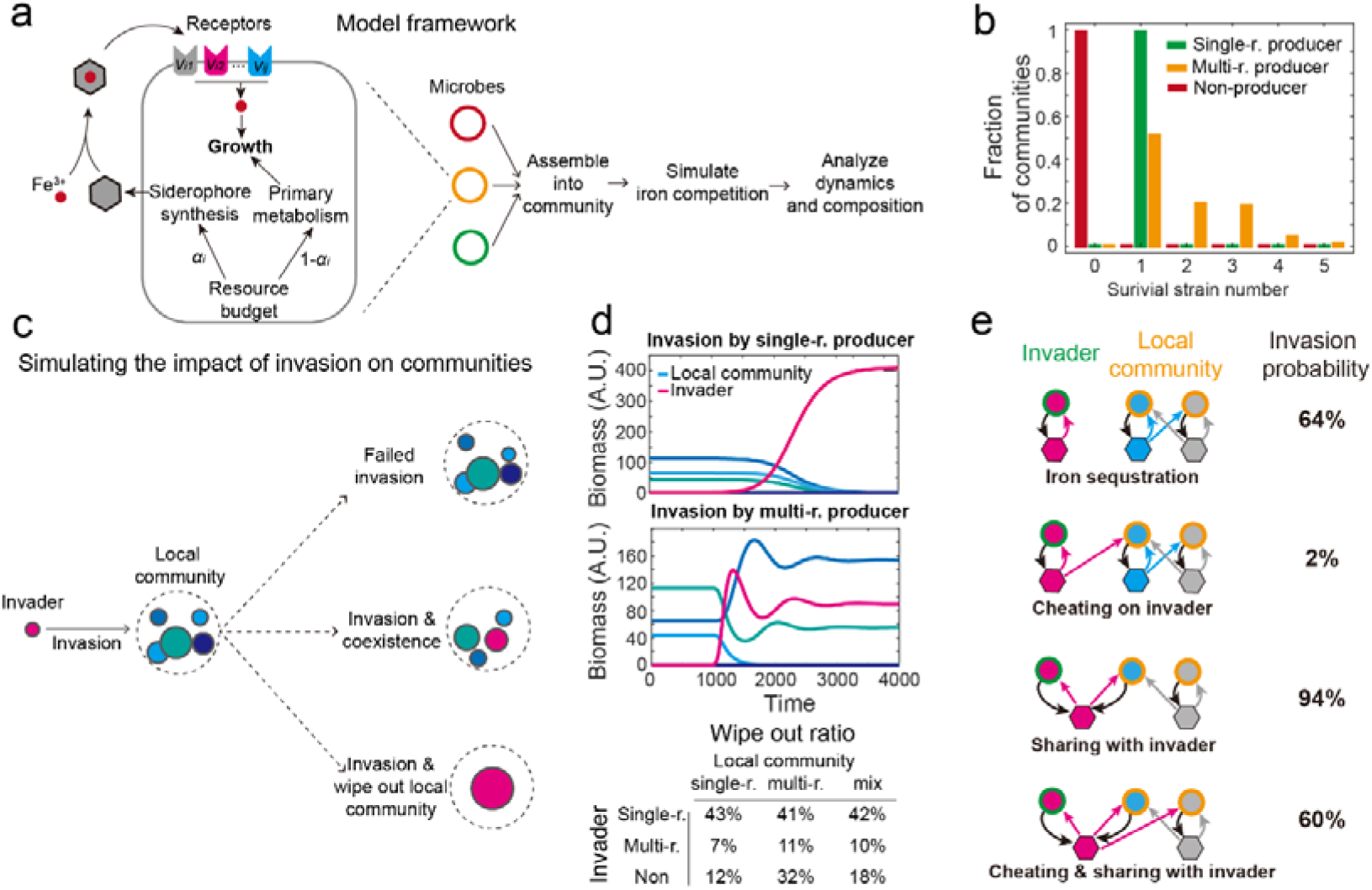
Mathematical model exploring the relationship between pyoverdine utilization strategies and community dynamics. **a.** Schematic diagram of the ecological iron competition model. Each microbe i can produce siderophores of type j with resource budget a_i_, and obtain iron by absorbing siderophore-iron complexes through corresponding receptors (fraction of receptors denoted as v_ij_). Growth rate is proportional to the total amount of absorbed iron and the fraction of resources allocated to primary metabolism, 1- a_i_. Various strains with distinct strategies were considered, including single-receptor producers (green), multi-receptor producers (orange), and non-producers (red), differing in their a_ij_ and v_ij_ values. 20 strains are assembled into communities, with diverse initial inoculation densities. They then compete in a chemostat-like model until reaching a steady state. **b.** Distribution of the number of species at steady state, for communities initiated with a single strategy type: single-receptor producers (green), multi-receptor producers (yellow), and non-producers (red). **c.** Schematic diagram depicting the invasion scenario together with the different possible outcomes. Putative invaders are introduced at low frequency into a local community at steady state. **d.** Examples illustrating common invasion dynamics. Top panel: a local community with three species (various blue shades) is invaded by a single-receptor producer (purple). Middle panel: a local community is invaded by a multi-receptor producer. Bottom panel: summary statistics showing the proportion of cases, in which invaders completely wiped out the local community as a function of their pyoverdine utilization strategy. **e.** The likelihood of a single-receptor producer to invade a local community of multi-receptor producers for four different iron interaction networks. Scenarios from top to bottom show: (1) no overlap in siderophore utilization between the local community and the invader; (2) multi-receptor producer from the local community can cheat on invader’s siderophore; (3) invader and one multi-receptor producer share the same siderophore; (4) scenario combining (2) and (3). Schematics are simplifications and do not reflect the actual number of species in the local community.

To delve deeper into the connection between pathogenicity and pyoverdine utilization strategies, we simulated the consequences of invasion by introducing a strain (“invader”) with a higher resource budget (available for intrinsic growth and siderophore production) into a steady-state community (“local community”) (Figure 5c). Across numerous parameters sets, we consistently observed that invasions by single-receptor producers and non-producers tended to disrupt local communities (Figure 5d). Successful single-receptor producers typically wiped out the local community. Conversely, successful multi-receptor producers integrated themselves into and co-existed with the local community (Figure 5d). In brief, the more severe consequences of invasion provide a plausible explanation for the association between certain strategies and pathogenicity.

Invasion probability further depended on the pyoverdine lock-key relationship between the invader (here modelled as a single-receptor producer, *e.g.,* matching a pathogenic lifestyle) and the local community (Figure 5e). The probability of successful invasion was close to zero if the local community could cheat on the invader’s siderophore, which is more likely when the invader produces siderophores with low Syn/Rec ratio. Conversely, invasion probability was high if the invader could produce an exclusive siderophore, or if one of the local community members produced the same siderophore as the invader (*i.e.*, enabling cheating of the invader). Taken together, our modelling results reveal that the relative proportion of single- and multi-receptor producers and their interactions via siderophores has fundamental consequences for community diversity and network complexity.

## Discussion

Predicting interactions between microbes from sequence data offers exciting opportunities for understanding microbiome assembly and stability, and may lay the foundation of biotechnological and medical microbiome interventions. While sequence-to-interaction mapping has predominantly been carried out for primary metabolism involving resource consumption, conversion, and cross-feeding, there are few approaches to reconstruct microbial interactions based on secondary metabolites (antibiotics, toxins, siderophores, surfactants) ^38-40^. In our paper, we offer such an approach by developing a co-evolution inspired computational approach to infer iron interaction networks mediated by pyoverdines (a class of iron-scavenging siderophores) within communities of *Pseudomonas* bacteria. Pyoverdines can both promote (through molecule sharing) and inhibit (through iron blocking) the growth of other strains, depending on a molecular lock-key (receptor-pyoverdine) mechanism^23,29^. Our co-evolution pairing algorithm managed to pair 188 pyoverdines types and 4547 receptors into 47 lock-key groups. Our experimentally validated approach allowed us to reconstruct the iron interaction network of 1928 *Pseudomonas* strain. We found intriguing network differences between habitats (soil, plant, water, human-derived habitats) and between microbial lifestyles (pathogenic and non-pathogenic). Large and highly connected siderophore-mediated iron interaction networks occurred among non-pathogenic environmental strain, whereas small and disconnected network dominated among pathogenic strains. These results suggest that selection pressures shaping bacterial interaction networks differ fundamentally between habitats and lifestyles.

Our sequence-to-ecology approach underscores several challenges associated with the reconstruction of interactions driven by secondary metabolites. The first challenge is that the chemical structures of secondary metabolites are often difficult to infer from sequencing data because the metabolites are produced by non-ribosomal peptide and polyketide synthesis pathways. The second challenge is to identify pyoverdine receptor genes among the many different types of siderophore receptor genes each strain possesses. We solved these challenges in a previous paper^31^, where we developed approaches based on feature sequences that allowed us to infer the chemical structure of 188 pyoverdines and identify 4547 pyoverdine receptor genes. The third challenge was to pair pyoverdines to matching receptors within and across strains. While co-evolution analyses are a widely used computational tool, employed in diverse areas ranging from ab initio protein structure to host-pathogen interaction predictions ^41^, we could not use existing algorithms, such as DCA, SCA, and Evoformer ^42-44^. The reason is that these classical site-based co-evolution methods depend on paired sequences between which the degree of covariation is quantified, yet the existence of multi-receptor producers impeded direct assignments of synthetase-receptor pairs. For this purpose, we developed an unsupervised learning algorithm (called Co-evolution Paring Algorithm), which yielded 47 receptor-synthetase lock-key pairs. This step was essential to reconstruct iron interaction networks. Our new pipeline has the potential to be applied to several other microbial traits. For example, microbial membrane receptors co-evolve with phages^45,46^, and pairing phages with the receptor they utilize for infection could provide insights into host-pathogen co-evolution and thus bacteria-host interaction networks in natural communities.

Our sequence-to-interaction mapping together with the mathematical model yielded several new biological insights into the ecology and evolution of microbial iron interaction networks. First, multi-receptor producers seem to be the glue of iron interaction networks. They harbor a large diversity of pyoverdine and receptor types, which can connect many other single- and multi-receptor producers and foster the formation of large and highly connected networks. Second, networks of non-pathogenic species and communities in natural soil and water habitats were large and highly interconnected. These networks were dominated by multi-receptor producers and were more open to invasion. In other words, new strains could easily integrate themselves into existing communities without much disturbance to the local community. This suggests that selection for siderophore and receptor diversity is particularly high in species-rich habitats. Third and in contrast to the above, networks of pathogenic species and communities in human-derived habitats were small, scattered and had low complexity. They were dominated by single-receptor producers with a compromised repertoire of pyoverdine and receptor types. Single-receptor producers were predicted to be the invaders with severe consequence, having a high potential to disrupt and displace the local community. Utilization of exclusive siderophore groups increases the chances of successful invasion. These results suggest that selection favors exclusive siderophores in pathogens as successful invasion is a key aspect of their lifestyle. Fourth, strains with a strict cheating strategy (non-producers) occur but are relatively rare (13.6%). However, it is important to note that our approach yields a conservative non-producer estimate because regulatory non-producers that have a synthetase cluster but do not express it also occur^22^ but cannot be detected with our approach. Given that non-producers are fully dependent on siderophore producers, it is intuitive to understand that they are (i) more common in environmental habitats featuring many multi-receptor producers with a rich pyoverdine repertoire, and (ii) best at invading multi-receptor communities.

Our insights on pyoverdine utilization strategies and their consequences for community dynamics reveal ways of how siderophore-mediated interactions could be leveraged for biotechnological applications ^47^. In this context, there is great interest in using probiotic strains in agriculture to protect crops from infections by bacterial plant pathogens. There is increasing evidence that siderophore-mediated interactions play a key role in this process^29,30^. For example, it was shown that plant-beneficial *Chryseobacterium* strains use their siderophores to suppress the plant-pathogen *Ralstonia solanacearum* ^48^. The approach has recently been extended to human pathogens, which were found to be suppressed by exclusive siderophores from environmental *Pseudomomnas* spp. ^47^. While these studies explored siderophore-interactions across the species boundaries without clear knowledge on the specific receptor setup, we here propose a strategic two-pronged design approach. First, single-receptor producers with exclusive siderophores should be designed and used as probiotics to competitively exclude pathogens in simple communities. Second, multi-receptor producers with an exclusive siderophore against the pathogen and cheating receptors utilizing the pathogen’s siderophores should be designed and used as probiotics in more complex communities. They could integrate themselves without major disruption of the local community, yet still competitively exclude the pathogen via the exclusive siderophore.

In conclusion, we succeeded in developing a sequence-to-interaction mapping approach for siderophores that has a high potential to deliver new insights into microbial ecology. Given that iron is a key trace element that is limited in most environments, siderophore-mediated interactions are an ideal entry point for secondary metabolite analysis from sequence data. While we focused on *Pseudomonas* strains, we know that siderophore-mediated interactions occur across the species boundaries. For example. *P. aeruginosa* possesses receptors to take up enterobactin produced by *Enterobacteriaceae* spp. and schizokinen produced by. *Ralstonia solanacearum* ^49^. Thus, the next step would be to apply our concepts to more diverse bacterial communities to derive microbiome-level iron-interaction maps that could further guide rational designs for biotechnological microbiota interventions.

## Data Availability

The source code and parameters used are available in the supplementary material.

## Acknowledgements

We thank Xuedi Huang for the insight on pyoverdine receptor analysis.

## Author contributions

Shaohua Gu performed the majority of computational and experimental data analysis in this research and drafted the manuscript. Zhengying Shao and Shenyue Zhu performed the experiment of testing pyoverdine-mediated interaction between *Pseudomonas* strains. Zeyang Qu performed the mathematical model exploring the relationship between pyoverdine utilization strategies and community dynamics. Di Zhang developed unsupervised co-evolution pairing algorithm for identifying self-receptor and Richard Allen offered the insight on this algorithm. Yuanzhe Shao, Ruolin He, Jiqi Shao, Guanyue Xiong assisted in cleaning up the codes. Alexandre Jousset and Ville-Petri Friman offered insightful comments and assisted in revising and writing of the manuscript. Rolf Kümmerli and Zhong Wei oversaw the project, designed experiments and revised the manuscript. Zhiyuan Li conceptualized, oversaw the project and revised the manuscript.

