## Supplementary_Information for "Siderophore synthetase-receptor gene coevolution reveals habitat- and pathogen-specific bacterial iron interaction networks"

Shaohua Gu *et al.*

*Corresponding authors

Zhong Wei.

Rolf Kümmerli.

Zhiyuan Li.

**This file includes:**

**Methods 1-10**

**Supplementary Figure S1-S7**

**Supplementary Table S1-2**

**Method**

1. **Construction of phylogeny tree**

The phylogenetic tree depicted in Figure 1a was constructed utilizing the PhyloPhlAn3 pipeline^1^. PhyloPhlAn is a comprehensive pipeline that encompasses the identification of phylogenetic markers, multiple sequence alignment, and the inference of phylogenetic trees. In this analysis, we employed over 400 universal genes defined by PhyloPhlAn as our selected phylogeny markers. Subsequently, the taxonomic cladogram was generated using the iTOL web tool ([http://huttenhower.sph.harvard.edu/galaxy/](https://huttenhower.sph.harvard.edu/galaxy)).

**2. Utilizing the co-evolution relationship between synthetases and receptors to identify self-receptors in producers**

To establish all lock-key relationships between synthetases and receptors in their sequence spaces (See Section 3 for results), it is necessary to identify the self-receptors in each producer strain. Assuming that the strongest co-evolution occurs between the synthetase and its cognate self-receptor, the pipeline of identifying self-receptors consists of following three key parts:

**Part 1. Calculation of the indel distance matrix between pyoverdine synthetase sequences**

To accurately quantify the evolutionary distance between synthetases, a more accurate method than the full sequence alignment is required. In the case of module- or domain-level duplication, deletion, or insertion event, the p-distance between two closely related sequences can become drastically high, even for single-receptor producers whose receptor sequences belong to the same classification group. This phenomenon is common and can potentially lead to the erroneous clustering of synthetase genes. Consequently, we undertook a two-step approach to enhance the accuracy of calculating sequence distances between synthetase genes.

**Step 1.** To address this issue, we initially conducted a global sequence alignment between any two synthetase feature sequences employing the Needleman-Wunsch algorithm. Utilizing the BLOSUM50 scoring matrix, we categorized all loci as “matched”, “similar”, or “unmatched”, based on their sequence similarity. Subsequently, we eliminated consecutive unmatched loci that extended for more than $L_{unmatch}$ amino acids, as these were deemed fragment mismatches resulting from module- or domain-level indels. The primary objective of this step was to mitigate the sequence distance between synthetase sequences belonging to strains within the same group. In our algorithm, the threshold $L_{unmatch}$ was set at 10 amino acids.

**Step 2.** For the remaining sequence segment, the ultimate distance, denoted as ${Distance}_{syn}$, was determined as:

$${Distance}_{syn}=pdistance*(1-p_{match})$$

Here, $pdistance$ represents the p-distance of the remaining sequence, while $p_{match}$ signifies the proportion of consecutive matched loci exceeding $L_{match}$ amino acids in length. Given the fewer consecutive matched loci observed between strains belonging to different groups, this step was implemented to further accentuate the disparities in synthetase sequence distances between strains within the same group and those in different groups. In our algorithm, we set the threshold for $L_{match}$ at 10 amino acids.

**Part 2. Calculation of the correlation between synthetase and receptor sequence distance matrices**

To effectively quantify the co-evolutionary relationship between synthetase and receptor sequences, we used the correlation between the distance matrices of synthetic and receptor sequences. We determined self-receptor pairs by checking if the grouping of synthetase sequences matched the grouping of receptor sequences. Recognizing that the Pearson correlation coefficients between synthetase and receptor sequence distance matrices could be sensitive to minor changes in sequence distances, we executed the following two steps to robustly quantify the correlation between synthetase and receptor sequence distance matrices:

**Step 1.** we applied binarization to both the synthetase and receptor sequence distance matrices using respective thresholds, $T_{syn}$ and $T_{rec}$. The distance matrix for synthetase was the indel distance matrix described in the preceding Part 1, and the distance matrix for receptors was calculated by alignment of the FpvA feature sequence (cite the method paper). Elements exceeding the threshold were assigned a value of 1, while elements falling below the threshold were assigned a value of 0.

**Step 2.** Following binarization, it's possible that indirect connections could exist, involving intermediate strains, in addition to the direct connections between strains. In order to distinctly separate multiple groups, we connected all connected components that are linked to each other, directly or indirectly, as one group. Then, the distance between strains belonging to different groups was standardized to 1, while the distance between strains within the same group was consistently set to 0. This connected component approach created an unweighted network encompassing all strains. This network preserved the connection information between strains while effectively mitigating any disturbances caused by minor fluctuations in sequence distances. In our algorithm, we designated both $T_{syn}$ and $T_{rec}$ as 0.3.

**Part 3. Unsupervised Co-evolution Pairing Algorithm for identifying self-receptor**

Using the correlation calculated in Part 2, We developed an unsupervised algorithm that leverages random sampling and simulated annealing to identify the self-receptor from a multitude of receptors in each multi-receptor producer. An overview of our algorithm is presented as follows (as depicted in Figure 3a):

**Step 1.** Initial processing.

1) In the case of $N$ multi-receptor producer strains, we initiated the process by computing the synthetase indel distances between all pairwise strains, as described in Part 1. This yields a synthetase sequence distance matrix with dimensions $N\times N$*.*

2) We randomly chose one receptor for each strain to construct the initial receptor list. Subsequently, we calculate the receptor sequence distance matrix, which is also of size $N\times N$, based on this initial receptor list.

3) We executed the connected component clustering procedure for both the synthetase and receptor sequence matrices, as described in Part 2. Following this, we computed the Pearson correlation coefficient between these two matrices.

**Step 2.** Random sampling.

We randomly selected $N_{batch}$ strains and introduced random perturbations to the receptor numbers associated with these strains, thus generating a perturbed receptor list.

**Step 3.** Calculation of the correlation coefficient.

1) We recalculated the receptor sequence distance matrix, by the perturbed receptor list.

2) The connected component clustering procedure in Part 2 was applied to the receptor distance matrix. the correlation coefficient between the synthetase and receptor distance matrices was calculated by methods in Part 2.

**Step 4.** Simulated annealing

According to the correlation coefficient calculated in Step 3:

1) If the correlation coefficient is smaller than the previous value, we would accept the perturbed receptor list.

2) If the correlation coefficient failed to decrease, we would accept the change of the receptor list with a probability denoted as $P_{accept}$.

**Step 5.** We returned to Step 2 and continued the interaction, until the correlation coefficients converge or a maximial number of iterations is reached.

In each iteration, the number of randomly selected strains was adaptive. We employed a smaller $N_{batch}$ when the number of iterations was limited, aiming to achieve a faster rate of correlation coefficient improvement. Conversely, when the number of iterations was extensive, we opted for a relatively larger $N_{batch}$ to introduce a wider range of perturbations. In our algorithm, the value for $N_{batch}$ was set as follows:

$$N_{batch}=\left\{ \begin{aligned} Random integer from \left[ 1, 5 \right], iterations<20000 \\ Random integer from \left[ 1, 10 \right], iterations>20000 \end{aligned} \right.$$

Furthermore, the inclusion of the simulated annealing step was instrumental in preventing the algorithm from getting stuck in local optimal solutions. In our algorithm, the acceptance probability, denoted as$P_{accept}$, was configured as follows:

$$P_{accept}=exp(\frac{corr^{'}-corr}{1/S_{iteration}})$$

Here, in the acceptance probability calculation, $corr^{'}$ represents the correlation coefficient derived from the perturbed receptor list, $corr$ denotes the correlation coefficient from the original receptor list, and $S_{iteration}$ signifies the current iteration count.

**3. Co-evolution Pairing Algorithm**

Among all multi-receptor producers, there are 678 synthetases and 2812 receptors in total. First, considering that NRPS pathways mainly evolve by large genetic rearrangement like recombination, we used the synthetase feature sequences (concatenated Amotif4-5 regions with consideration of recombination, See Method 3 for details) to build the 678x678 synthetase distance matrix (Figure 3a). We then picked a random receptor as putative self-receptor for each multi-receptor producer and used the receptor features sequences (168 Pro to 295 Ala) to calculate the corresponding 678x678 receptor distance matrix. Subsequently, we calculated co-evolution coefficient *cr*, defined as the Pearson's correlation coefficient between the two matrices (see Method for details). The initial random self-receptor assignment resulted in poor co-evolution coefficients. We thus introduced an iterative optimization process, during which putative self-receptors were shuffled within each multi-receptor producer. We discarded iterations that decreased *cr* values and continued with those that increased *cr* values until an optimization plateau was reached (Figure S1, *cr* = 0.84). We predicted the self-receptor of all multi-receptor producers based on the final assignment.

**4. DNA extract of *Pseudomonas* strains**

We used 24 *Pseudomonas* strains that were originally isolated from tomato rhizosphere^2^ to test the effects of pyoverdine on interactions between strains. The genomic DNA of *Pseudomonas* strains was initially extracted using Invitrogen PureLink® Genomic DNA kit. The DNA quantity and quality were tested by the NanoDrop ND-1000 Spectrophotometer (Thermo Fisher Scientific). The DNA was purified further using the Quick-DNA Miniprep Plus kit.

**5. Illumina HiSeq sequencing**
For Illumina pair-end sequencing of each strain, at least 3μg genomic DNA was used for sequencing library construction. Paired-end libraries with insert sizes of ~400bp were prepared following Illumina’s standard genomic DNA library preparation procedure.Purified genomic DNA is sheared into smaller fragments with a desired size by Covaris, and blunt ends are generated by using T4 DNA polymerase. After adding an ‘A’ base to the 3' end of the blunt phosphorylated DNA fragments, adapters are ligated to the ends of the DNA fragments. The desired fragments can be purified through gel-electrophoresis, then selectively enriched and amplified by PCR. The index tag could be introduced into the adapter at the PCR stage as appropriate and we did a library quality test. At last, the qualified Illumina pair-end library would be used for Illumina NovaSeq 6000 sequencing (150bp*2, Shanghai BIOZERON Co., Ltd).

**6. PacBio Sequencing**

The whole genome sequencing was performed there by Pacific Biosciences Sequel II technology (PacBio). The DNA was made into SMRTbell libraries using the Express Template Prep Kit 2.0 from PacBio according to the manufacturer’s protocol. Samples were pooled into a single multiplexed library and size was selected using Sage Sciences’ BluePippin, which uses the 0.75% DF Marker S1 High-Pass 6 kb–10 kb v3 run protocol and S1 marker. A size selection cutoff of 8000 (BPstart value) was used. The size selected SMRTbell library was annealed and bound according to the SMRT Link Set Up and sequenced on a Sequel II.

**7. Genome Assembly**

The raw paired end reads were trimmed and quality controlled by Trimmomatic (version 0.36, <http://www.usadellab.org/cms/uploads/supplementary/Trimmomatic>) with parameters (SLIDINGWINDOW:4:15 MINLEN:75). Clean data obtained by above quality control processes were used to do further analysis.

Raw PacBio reads were converted to fasta format with Samtools Fasta (http://www.htslib.org/doc/samtools.html). The Illumina data was used to evaluate the complexity of the genome and correct the PacBio long reads. First, we used unicycler (https://github.com/rrwick/Unicycler) to perform genome assembly with default parameters and received the optimal results of the assembly. GC depth and genome size information were calculated by custom perl scripts which can help us to judge whether DNA sample contained contaminated or not. Finally, strain genome was circularized with Circlator (http://sanger-pathogens.github.io/circlator/).

**8. Genome Annotation**

For the prokaryotic organism, we used ab initio prediction method to get gene models for strain xx. Gene models were identified using GeneMark. Then all gene models were blastp against non-redundant (NR in NCBI) database, SwissProt (http://uniprot.org), KEGG (<http://www.genome.jp/kegg/>), and COG (<http://www.ncbi.nlm.nih.gov/COG>) to do functional annotation by blastp module. In addition, tRNA were identified using the tRNAscan-SE (v1.23, http://lowelab.ucsc.edu/tRNAscan-SE) and rRNA were determined using the RNAmmer (v1.2, http://www.cbs.dtu.dk/services/RNAmmer/).

**9. Measuring the growth of Pseudomonas strains and their pyoverdine production**

All *Pseudomonas* strains were stored at −80°C. Prior to the experiments, a single colony of each strain was selected randomly, grown overnight in lysogenic broth (LB), washed three times in 0.85% NaCl, and adjusted to an optical density at 600 nm (OD_600_) of 0.5 using a spectrophotometer (Spectra Max M5, Sunnyvale, CA, USA). To quantify the growth and siderophore production of each pseudomonad strains under iron-limited conditions, we transferred 2 mL of overnight cultures into new 250 mL glass flask containing 150 mL MKB medium (K_2_HPO_4_ 2.5 g L^-1^, MgSO_4_·7H_2_O 2.5 g L^-1^, glycerin 15 ml L^-1^, casamino acids 5.0 g L^-1^, pH 7.2) in three-fold replication. Following 24 h of incubation at 30°C with shaking (rotary shaker set at 170 rpm), we measured growth (OD_600_) with a spectrophotometer at room temperature (SpectraMax M5, Sunnyvale, CA, USA) and then harvested the cell-free supernatant from bacterial cultures by centrifugation (8,000 rpm, 8 min at 4°C) and filtration (using a 0.22 µm filter). The supernatant was then divided into two parts for i) measuring pyoverdine production and ii) testing the effects of pyoverdine on interactions between Pseudomonas strains. Briefly, pyoverdine production levels were measured (relative fluorescence units (RFU) with excitation: 400 nm and emission: 460 nm) with a spectrophotometer at room temperature (SpectraMax M5, Sunnyvale, CA, USA).

**10. Testing the effects of pyoverdine on interactions between Pseudomonas strains**

To avoid interference from other metabolites as much as possible, we adapted the method of Butaitė et al. ^3^ to crudely purify pyoverdine from the supernatants of 20 producers collected through the above steps. For the cross-feeding assay, we suspended each purified pyoverdine in 2 mL Milli-Q water and passed the solution through a 0.22 µm filter.

Following the above steps to obtain 24 strains of bacterial fluid (OD_600_ = 0.5), we diluted invader pre-cultures 100-fold into new 96-well plates and subjected them to the following three experimental conditions in three-fold replication.  (1)${SN}_{limited}$: each strain individually growing in 180 µl 10% MKB medium supplemented with 20 µl aqueous solution of pyoverdine crude extract. (2) ${SN}_{replenished}$: each strain individually growing in 180 µl iron rich 10% MKB medium supplemented with 20 µl aqueous solution of pyoverdine crude extract (removes the effect of pyoverdine but retains the effect of other metabolites). The iron-rich condition was achieved by adding iron (III) solution (1 mM FeCl_3_·6H_2_O, 10 mM HCl) into MKB medium (final concentration equaling 50 μM). (3) ${SN}_{control}$: each strain individually growing in 180 µl iron-limited 10% MKB medium supplemented with 20 µl of 0.85% (w/v) NaCl instead of supernatant (control mimicking the addition of spent medium). We measured each pseudomonad strain growth [OD_600_] of each replicate after 24 h of incubation at 30°C under static conditions.

Subsequently, we calculated the effect of each producer's pyoverdine crude extract on each pseudomonad strain growth as growth effect, denoted as ${GE}_{treatment}$, was determined as:

$${GE}_{treatment}=\left( \left( {SN}_{treatment}/{SN}_{control} \right)-1 \right)*100$$

where ${SN}_{treatment}$ is ${SN}_{limited}$ or ${SN}_{replenished}$. For this calculation, we took the average effects across the three replicates. From these measures, the net GE of pyoverdine can be measured as ${GE}_{pyo}={GE}_{li}-{GE}_{re}$. This is possible because we used the exact same supernatants for ${SN}_{limited}$ and ${SN}_{replenished}$, but pyoverdines are only important for growth in the former and not in the latter condition, where iron is available in excess^2^. In principle, GE_Pyo_ > 0 indicates pyoverdine-mediated facilitation. However, because there is substantial experimental variation between experimental replicates, we increased a threshold value of GE_Pyo_ > 0.05 and classified values above this threshold as positive interactions, where the receiving strain can use the respective pyoverdine for iron acquisition (interaction type 1). Conversely, GE_Pyo_ ≤ 0.05 values were classified as neutral or negative interactions, where the receiving strain cannot use the respective pyoverdine for iron acquisition (interaction type 0).

**Supplementary Figures**

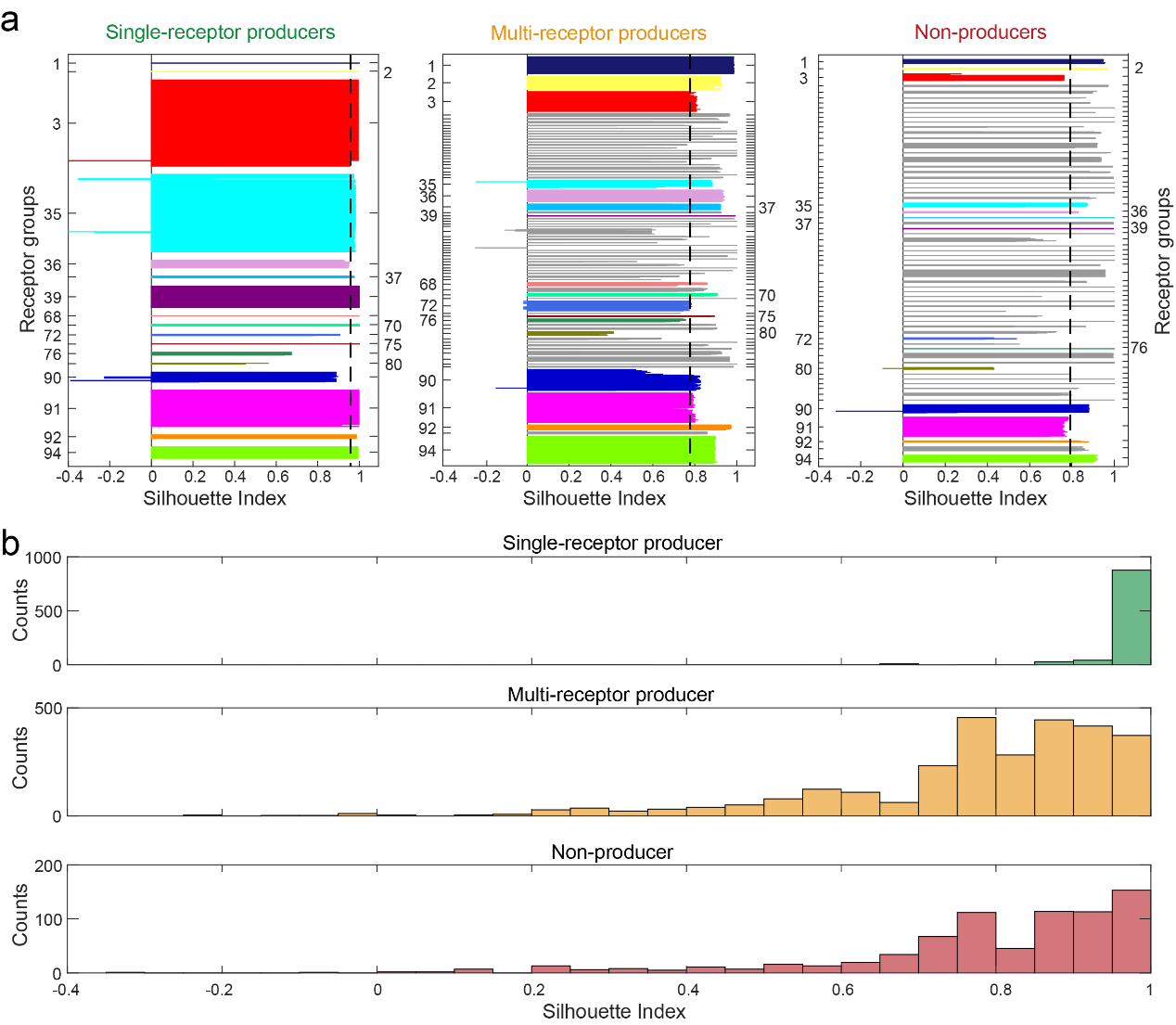

**Figure S1 Silhouette index analysis on the compactness of all receptor groups in single-receptor producers (left panel), multi-receptor producers (middle panel) and non-producers (right panel).** Colors in **a** represent all the 17 receptor groups found among single-receptor producers. All other receptor groups are shown in black. The dashed vertical lines represent the average of the Silhouette index across all the receptor groups within each strain class.

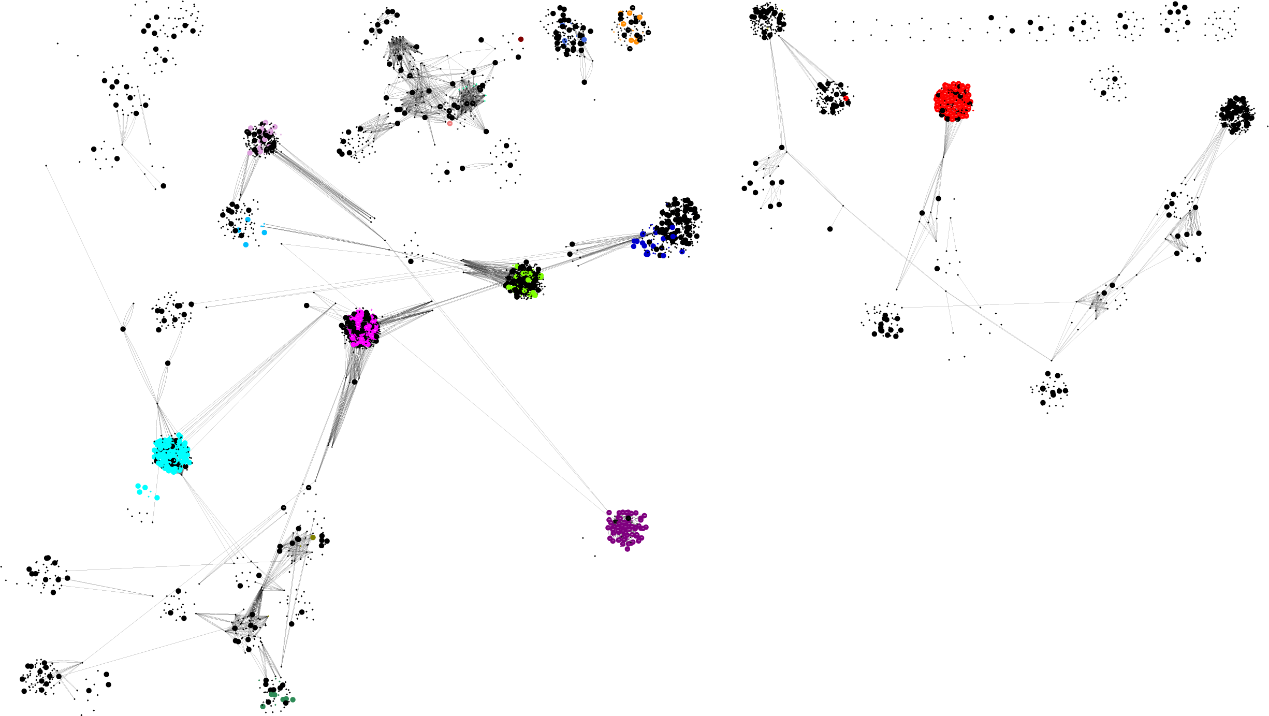
**Figure S2 The sequence similarity network of all 4547 FpvAs receptors built by Cytoscape's edge-weighted Prefuse force-directed layout.** Each FpvA is represented by a network node. Colors indicate the 17 FpvA groups found among the single-receptor producers (same color code as Figure 1c). FpvA receptors of non-producers (small dots) and multi-receptor producers (large dots) are colored gray and black, respectively. The width of an edge represents the sequence similarity between connected nodes. Edges connecting receptors within the same group were hidden. Only edges with a similarity higher than 50% and less than 70% to other groups were displayed for each group.

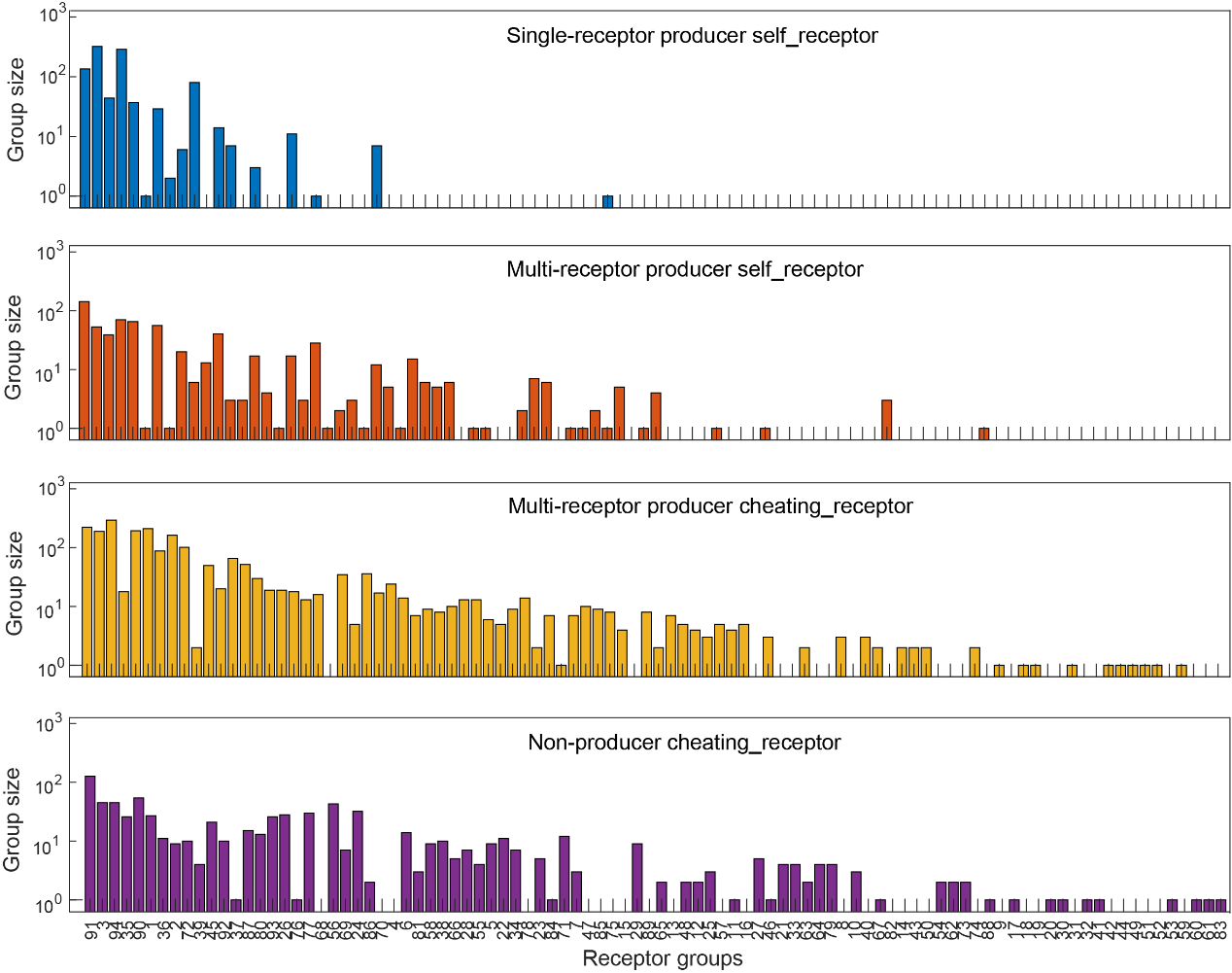

### Figure S3 The 94 FpvA receptor groups (sorted by group size) and their frequency among single-receptor producer self_receptor, multi-receptor producer self_receptor, multi-receptor producer cheating_receptor (non-self-receptor) and non-producer cheating_receptor.

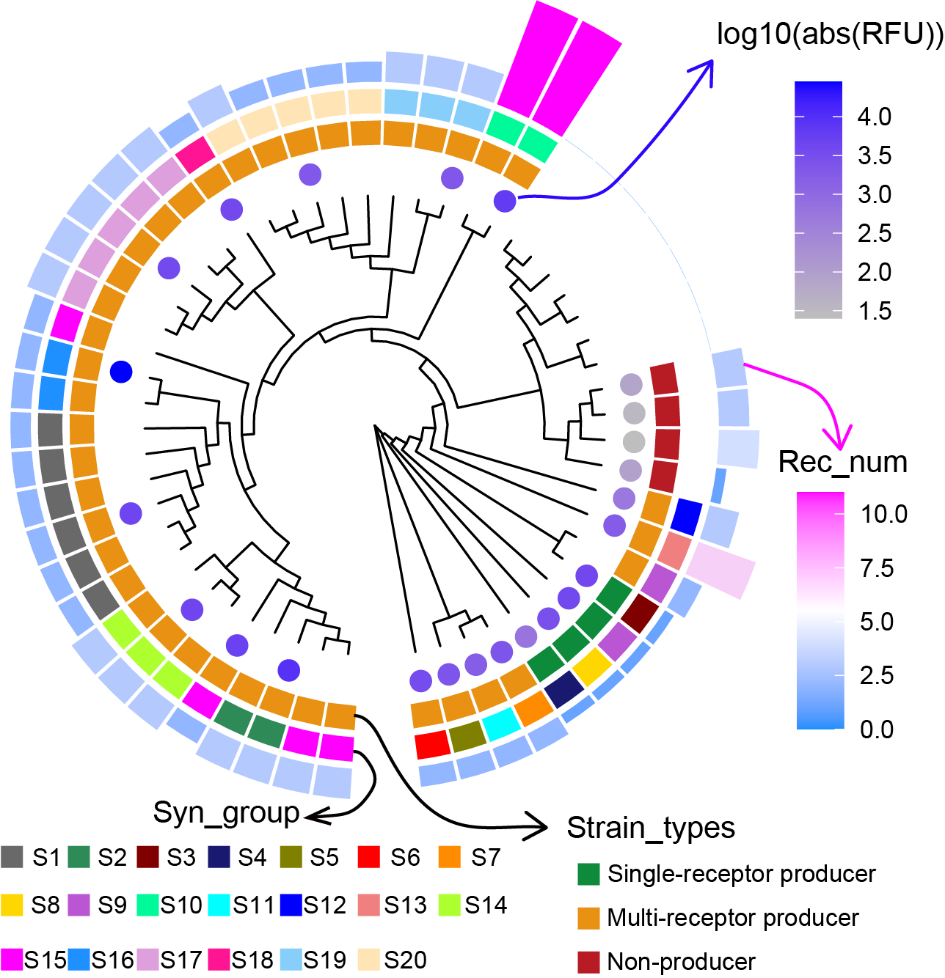

### Figure S4 Phylogeny tree of the 55 pseudomonas strains, based on concatenated alignments of 400 single-copy conserved genes. Color and bar height in the outmost ring indicate the number of FpvA receptors present in each strain. Colors in the second ring distinguish the 20 synthetase groups. Colors in the third ring highlight the three basic classes of iron-utilization strategies. To remove duplicates, we randomly selected one strain from each synthetase group to carry out the cross-feeding experiments. Colors in the innermost ring indicate the mean pyoverdine production (measured across three replicates) for the 24 experimental strains.

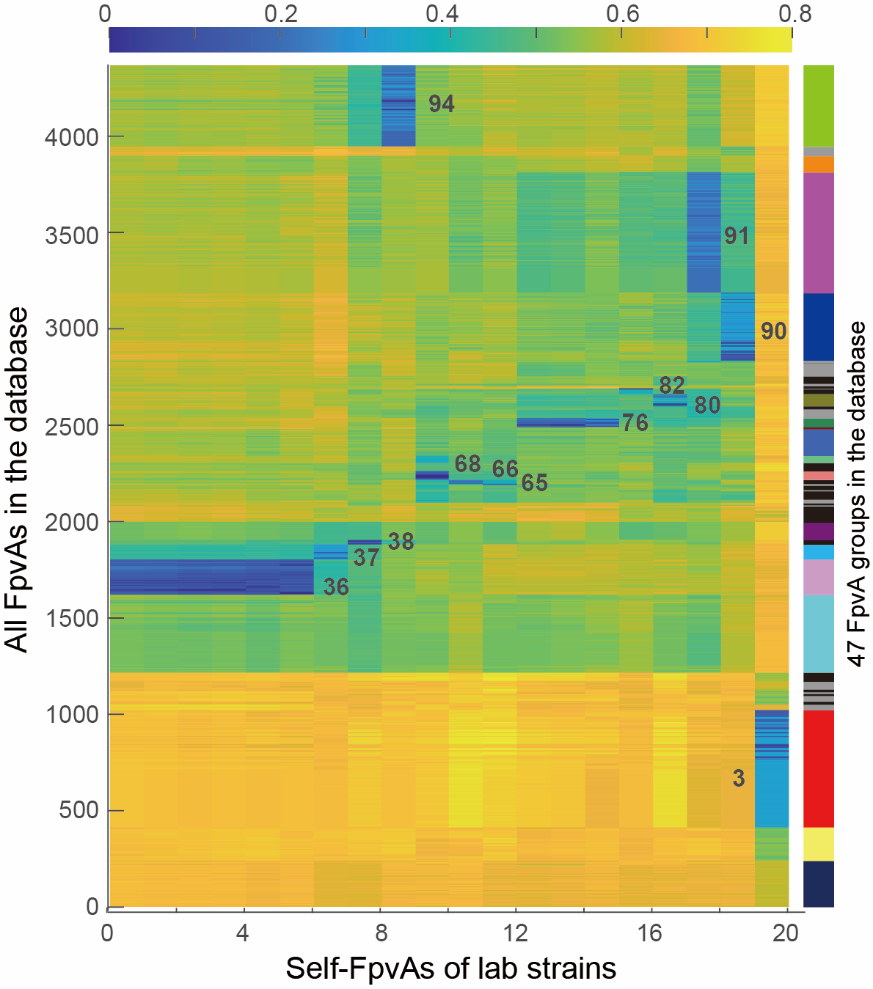

**Figure S5 Heatmap visualization of the feature sequence distances between the self-receptors of the 20 experimental producer strains and all receptors of the 47 lock-key groups in our database.** The self-receptors of the 20 experimental producer strains belong to 13 receptor groups, which are marked by black text.

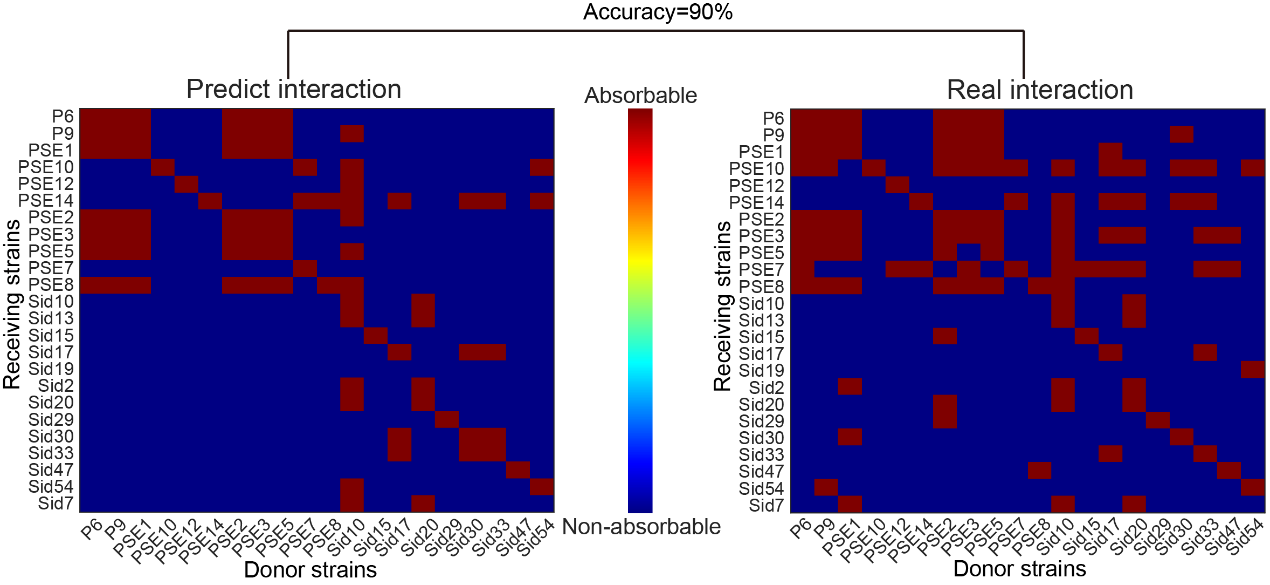
**Figure S6 Predicted pyoverdine mediated iron-interactions between laboratory strains based on genomic data mining and experimentally measured true pyoverdine-mediated iron interactions between strains.**

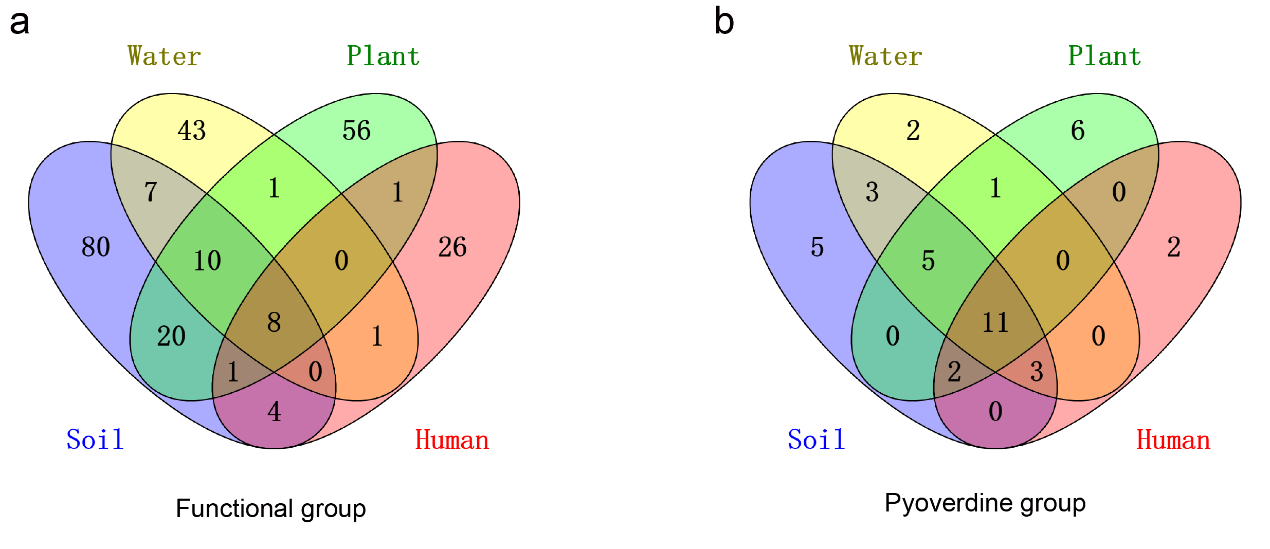

### Figure S7 Venn diagrams showing the overlap in siderophore functional groups (a) and pyoverdine lock-key groups (b) of Pseudomonas strains isolated from different habitats.

**Supplementary tables**

**Table S1 Distribution of *Pseudomonas* strain types in different habitats**

|  | Single-receptor producer (%) | | Multi-receptor producer (%) | Non-producer (%) |
| --- | --- | --- | --- | --- |
| Soil | | 27.48 | 56.87 | 15.65 |
| Plant | | 36.32 | 53.85 | 9.83 |
| Water | | 42.74 | 43.55 | 13.71 |
| Human | | 86.06 | 10.02 | 3.91 |

**Table S2 Pathogenic and non-pathogenic information for species with more than 5 strains**

| Species name | Strain number | Is pathogenic | Siderophore functional group number |
| --- | --- | --- | --- |
| *P.sp* | 621 | Nan | 34 |
| *P.aeruginosa* | 554 | Yes ^4^ | 3 |
| *P.fluorescens* | 135 | No ^5^ | 13 |
| *P.syringae* | 116 | Yes ^6^ | 5 |
| *P.putida* | 43 | No ^7,8^ | 18 |
| *P.protegens* | 23 | No ^9^ | 2 |
| *P.fragi* | 16 | Nan ^10^ | 0 |
| *P.stutzeri* | 16 | Yes ^11,12^ | 1 |
| *P.amygdali* | 15 | Yes ^13,14^ | 1 |
| *P.chlororaphis* | 15 | No ^15,16^ | 4 |
| *P.koreensis* | 15 | Nan ^17,18^ | 4 |
| *P.psychrotolerans* | 14 | Nan ^19,20^ | 1 |
| *P.brassicacearum* | 13 | Nan ^21^ | 4 |
| *P.coronafaciens* | 11 | Yes ^22^ | 1 |
| *P.oryzihabitans* | 11 | Yes ^23,24^ | 0 |
| *P.frederiksbergensis* | 10 | Nan ^25^ | 3 |
| *P.savastanoi* | 10 | Yes ^26^ | 1 |
| *P.moraviensis* | 9 | Nan ^27^ | 2 |
| *P.lundensis* | 8 | Nan ^28^ | 1 |
| *P.mandelii* | 8 | Nan ^29,30^ | 3 |
| *P.monteilii* | 8 | Nan ^31,32^ | 4 |
| *P.asiatica* | 7 | nan^33^ | 1 |
| *P.parafulva* | 7 | No ^34^ | 4 |
| *P.poae* | 7 | Nan ^35^ | 2 |
| *P.asplenii* | 6 | Yes ^27^ | 2 |
| *P.helleri* | 6 | Nan ^36^ | 1 |
| *P.mendocina* | 6 | Yes ^37^ | 0 |
| *P.nitroreducens* | 6 | Nan ^38^ | 0 |
| *P.synxantha* | 6 | No ^39^ | 2 |
